# Genome-wide Prediction of microRNAs in Zika virus Genomes Reveals Possible Interactions with Human Genes Involved in the Nervous System Development

**DOI:** 10.1101/070656

**Authors:** Juan Cristina, Natalia Echeverría, Fabiana Gambaro, Alvaro Fajardo, Pilar Moreno

**Affiliations:** Laboratorio de Virología Molecular, Centro de Investigaciones Nucleares, Facultad de Ciencias, Universidad de la República, Montevideo, Uruguay.

## Abstract

Zika virus (ZIKV) is a member of the family *Flaviviridae*. In 2015, ZIKV triggered a large epidemic in Brazil and spread across Latin America. In November of that year, the Brazilian Ministry of Health reported a 20-fold increase in cases of neonatal microcephaly, which corresponds geographically and temporally to the ZIKV outbreak. ZIKV was isolated from the brain tissue of a fetus diagnosed with microcephaly, and recent studies in mice models revealed that ZIKV infection may cause brain defects by influencing brain cell developments. Unfortunately, the mechanisms by which ZIKV alters neurophysiological development remain unknown. MicroRNAs (miRNAs) are small noncoding RNAs that regulate post-transcriptional gene expression by translational repression. In order to gain insight into the possible role of ZIKV-mediated miRNA signaling dysfunction in brain-tissue development, we computationally predicted new miRNAs encoded by the ZIKV genome and their effective hybridization with transcripts from human genes previously shown to be involved in microcephalia. The results of these studies suggest a possible role of these miRNAs on the expression of human genes associated with this disease. Besides, a new ZIKV miRNA was predicted in the 3’stem loop (3’ SL) of the 3’untranslated region (3’UTR) of the ZIKV genome, suggesting the role of the 3’UTR of flaviviruses as a source of miRNAs.

## Introduction

Zika virus (ZIKV) is a member of the family *Flaviviridae*, whose natural transmission cycle involves mainly vectors from the *Aedes* genus, while humans are occasional hosts [1]. Clinical manifestations of disease caused by ZIKV range from asymptomatic cases to an influenza-like syndrome associated to fever, headache, malaise and cutaneous rash [2]. ZIKV genome consists of a single-stranded positive sense RNA molecule of approximately 10,800 nt in length [3]. Although ZIKV enzootic activity was reported in diverse countries of Africa and Asia, few human cases were reported until 2007, when a large epidemic took place in Micronesia [4]. A large ZIKV outbreak occurred in French Polynesia during 2013–2014 and then spread to other Pacific islands [5]. In early 2015, a ZIKV epidemic outbreak took place in Brazil, currently estimated at 440,000– 1,300,000 cases [6]. In November 2015, the Brazilian Ministry of Health reported a 20-fold increase in cases of neonatal microcephaly, which corresponds geographically and temporally to the ZIKV outbreak [7]. Due to this global threat, World Health Organization (WHO) declared a public health emergency of international concern on February 1^st^, 2016 [8].

ZIKV was detected by electron microscopy and RT-qPCR in brains and amniotic fluid of microcephalic fetuses, strengthening the causal link between ZIKV and increased incidence of microcephaly [9, 10]. Furthermore, recent studies show that ZIKV can infect human iPSC-derived neural progenitor cells (NPCs) *in vitro*, resulting in dysregulation of cell-cycle-related pathways and increased cell death [11]. The development of animal models permits to begin to understand the underlying pathology of ZIKV infection [12, 13]. Modeling ZIKV infection in mice revealed direct effects of ZIKV on neural precursor cells development, including proliferation, differentiation and cell death, which may link ZIKV with the development of microcephaly [14]. Notably, genes associated with microcephaly were downregulated in ZIKV-infected mice brains [14] and NPCs [11]. Multiple approaches will be needed to understand the pathogenesis of ZIKV infection [12].

MicroRNAs (miRNAs) are small regulatory non-coding RNAs, ranging from 19 to 24 nucleotides in length, that post-transcriptionally regulate target gene expression by inhibiting the translation of mRNA transcripts or leading to their cleavage [15]. While encoded not only by cellular genomes but also by viral genomes [16], miRNAs play a vital role in diverse biological processes, including development, apoptosis, tumorogenesis, proliferation, etc. [17]. Viral miRNAs are mostly identified by traditional cloning from virus-infected cells [18], nevertheless, computational prediction and hybridization analysis are also applied to viral miRNA identification [19, 20]. To date, RNA virus-encoded miRNAs have been identified in Hepatitis C virus [21], Human Immunodeficiency virus (HIV) [22], Bovine Leukemia virus (BLV) [23], Middle East Respiratory Syndrome (MERS) coronavirus [24] Hepatitis A virus (HAV) [25] and Ebolavirus [26]. Interestingly, miRNAs have also been identified in other members of the family *Flaviviridae*, like West Nile virus (WNV) [27] or Dengue virus (DENV) [28]. Very recent *in silico* studies revealed the miRNA coding capacity of the ZIKV genome [29]. In order to get insight into the role of ZIKV genome in relation to miRNA coding, we computationally identified new potential miRNAs on the ZIKV genome along with their target genes. Our study may help to better understand host-pathogen interaction as well as to contribute to the development of new antiviral strategies against ZIKV infection.

## Materials and Methods

### Viral sequences and prediction of pre-miRNAs by an *ab initio* approach

In this study, we used complete genome sequences of ZIKV strain Natal-RGN (GenBank accession number: KU527068), isolated in 2015, from the brain tissue of a fetus diagnosed with microcephaly [10]. VMir program [30] was used to analyze the ZIKV strain sequences. VMir is an *ab initio* prediction program specifically designed to identify pre-miRNA in viral genomes. Using this approach, ZIKV sequences were analyzed for possible pre-miRNA hairpin structures, using highly stringent filtering parameters (minimum hairpin size of 60 nucleotides, maximum hairpin size of 120 nucleotides, minimum hairpin score of 160, minimum window count of 25). VMir scores were calculated according to Grundhoof [30].

### Confirmation of putative pre-miRNA sequences

To discriminate real pre-miRNAs from other hairpin structures (pseudo hairpins) we employed iMiRNA-SSF approach [31]. This approach represents an improvement in the performance for accurate identification of miRNA precursors by combining negative sets with different distributions [31].

### Identification of mature miRNA sequences

With the purpose of extracting mature miRNA: miRNA* duplexes from pre-miRNA hairpins, we employed MiRduplexSVM [32]. This approach uses a novel SVM-based methodology and takes into account several aspects of the biogenesis of miRNAs, whereby a duplex is formed before the mature molecule is selected [32].

### Conservation of predicted miRNA sequences among ZIKV strains

In order to observe the degree of conservation of the predicted pre-miRNA and mature miRNA sequences in Natal-NGR ZIKV strain and strains from all types isolated elsewhere, the sequences from all available and comparable ZIKV strains for which the complete genome sequences are known, were downloaded from the GeneBank database. Once downloaded, the sequences were aligned using the MUSCLE program [33]. Consensus sequences (i.e. the calculated most frequent nucleotide residue found at each position of the sequence alignment) were established using Entropy-One software from the HIV database (available at: http://www.hiv.lanl.gov/). For names and accession numbers of ZIKV strains included in these studies, see Supplementary Material Table 1.

**Table 1.**
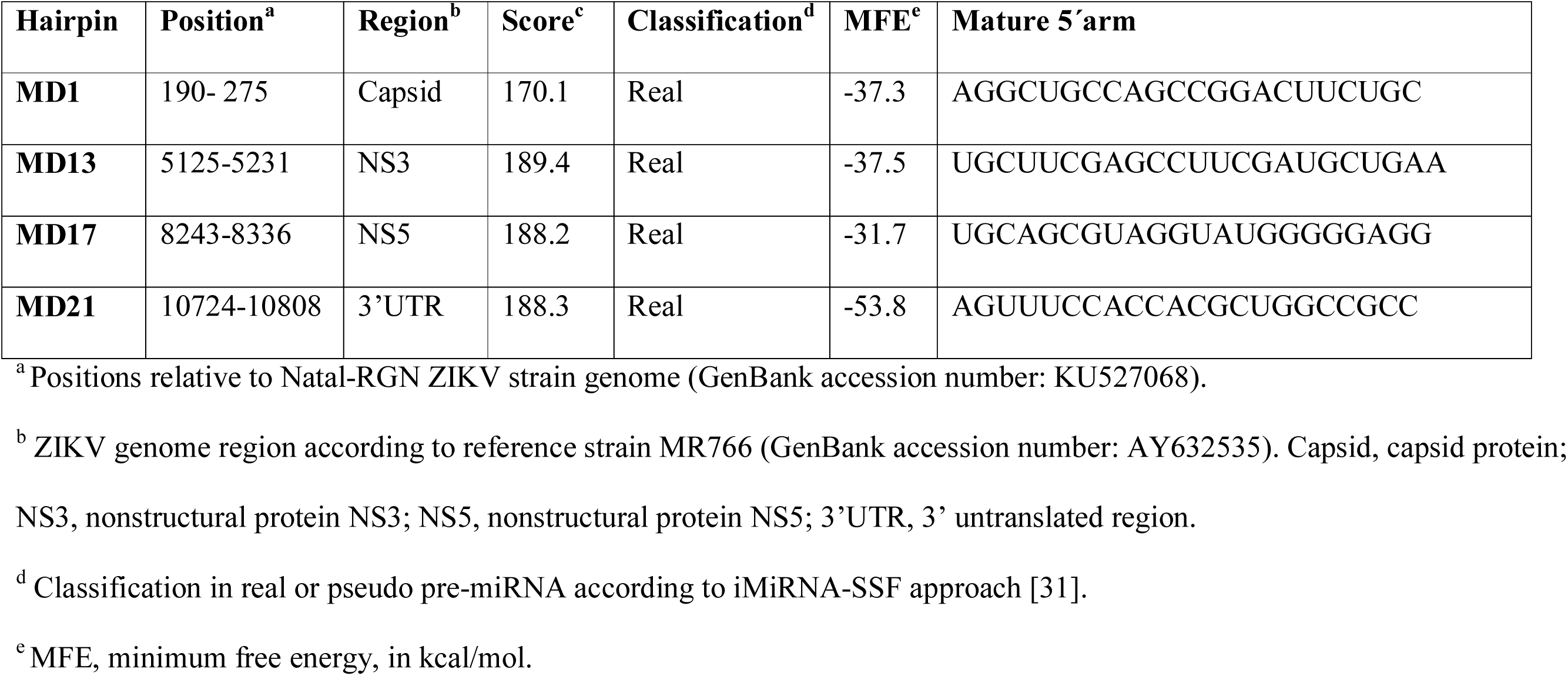
Predicted hairpin and mature sequences within Natal-RGN ZIKV genome.

### Prediction of secondary structure of miRNA precursors

The RNAfold web server [34] was used to predict the secondary structure of pre-miRNAs. Only default parameters were used.

### Prediction of potential gene targets

In order to identify the regulatory relationships between predicted miRNAs and their putative target gene transcripts we employed miRTar [35]. The parameters assigned for the hybridization were set to −14 kcal/mol of minimum free energy (MFE) for all hybridizations as a cut-off value and an alignment score ≥ 140. We studied the relation between predicted miRNAs and their target gene transcripts focusing on human genes involved in microcephaly. For names, accession numbers and their roles on brain development of the genes involved in these analyses, see Supplementary Material Table 2.

### Hybridization between gene transcript targets and mature ZIKV miRNA

In order to confirm an effective hybridization between gene transcript targets and predicted ZIKV miRNA, we employed the RNAhybrid [36]. RNA hybrid is a tool for finding the minimum free energy hybridization of a long and a short RNA and is widely used for miRNA target prediction. Pairing energy or minimum free energy (MFE) indicates the stability of the hybridization. A stringent condition was employed, with a MFE set at −30 kcal/mol for all hybridizations, as a cut-off value.

## Results

### Prediction of pre-miRNA stem-loop structures in Natal-RGN ZIKV genome

Computational prediction represents a widely used and effective strategy to identify novel miRNAs that can be further examined and validated by experimental approaches. In order to observe whether ZIKV genome could be folded into putative pre-miRNA stem-loop structures, we first analyzed its putative miRNA-encoding capacity using full-length sequences from Natal-RGN ZIKV genome (GenBank accession number: KU527068). We filtered VMir [30] output using stringent custom setting. The results of these studies revealed six pre-miRNA hairpins that were selected as potential hairpins and suggested the potential miRNA-coding capacity of the genome of this ZIKV strain. Then, in order to confirm the presence of these pre-miRNA structures, we classified them into real or pseudo pre-miRNAs by means of the iMiRNA-SSF approach [31]. Four out of the six pre-miRNA structures (named MD1, MD13, MD17 and MD21) were classified as real pre-miRNA. These structures are present in different genomic regions of the ZIKV genome (see Table 1).

### miRNAs sequence conservation among ZIKV strains genomes

As an RNA virus, ZIKV has a high degree of genetic variability and heterogeneity. Recent studies have estimated the rate of evolution of ZIKV from 0.98 to 1.06 × 10^−3^ substitutions per site per year (s/s/y) [37] and 1.20 × 10^−3^ s/s/y [38]. Besides, previous phylogenetic analysis characterized two major genetic lineages of ZIKV, the African and Asian lineages [39]. Very recently, phylogenetic analyses revealed the presence of three distinct lineages, (Asian/American lineage, African lineage 1 and African lineage 2) [40]. For these reasons, it is important to establish the degree of conservation of the pre-miRNAs and mature miRNAs sequences of predicted miRNAs in the genome of Natal-RGN ZIKV strain among ZIKV strains from all genetic lineages and isolated elsewhere.

In order to gain insight into this matter, all available and comparable ZIKV strains for which complete genome sequences are known were downloaded from the GenBank database and aligned using the MEGA6 program [41]. These strains were isolated elsewhere and represent all genetic lineages of ZIKV strains (for strains and accession numbers, see Supplementary Material Table 1). Once aligned, consensus sequences (i.e. the calculated most frequent nucleotide residue found at each position of the sequence alignment) were established and compared to predicted pre-miRNA and mature miRNA sequences in Natal-NGR ZIKV strain. The results of these studies revealed 98% identity among pre-miRNA MD1 and corresponding consensus sequences, and 100% identity among pre-miRNAs MD13, MD17 and MD21 and corresponding consensus sequences. 100% identity among mature miRNAs and consensus sequences were found in all cases. The secondary RNA structure of predicted pre-miRNA sequences and corresponding consensus sequences were identical in all cases (see Fig 1). The results of these studies revealed that these predicted pre-miRNAs and mature miRNAs sequences are highly conserved among ZIKV strains.

**Fig 1.**
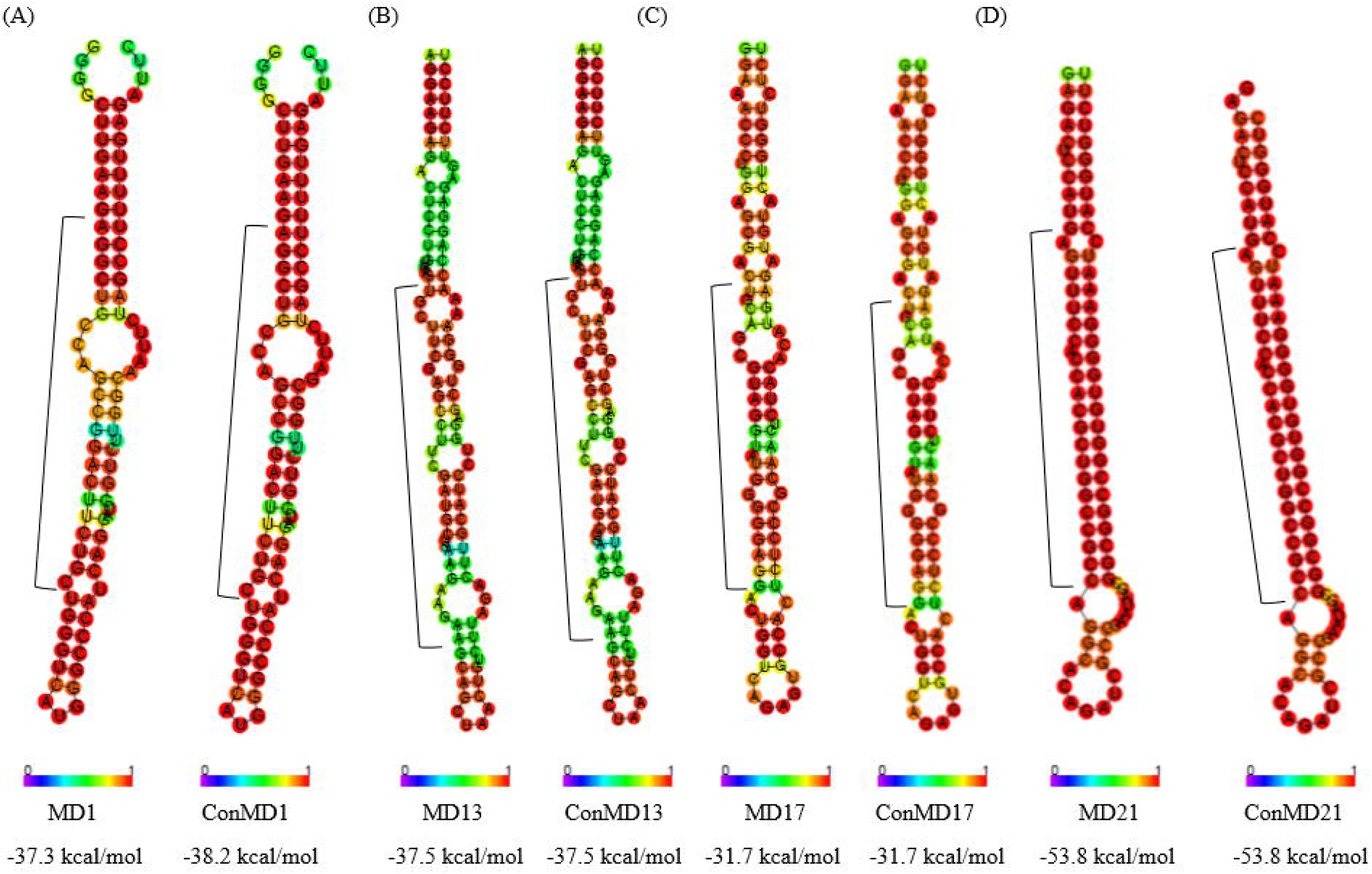
Predicted secondary structure of pre-miRNAs. In A to D, comparisons of the predicted secondary structure of the pre-miRNAs sequences found in Natal-NGR ZIKV strain and ZIKV corresponding consensus sequences are shown. Structures of the predicted pre-miRNA hairpins and their respective consensus (con) sequences are indicated by name at the bottom of the figure as well as the minimum free energy (MFE) values obtained for each structure. Bars at the bottom of the structures denote base pair probabilities. Only centroid structures are depicted. Mature miRNAs sequences are indicated by a square bracket next to each structure.

### Prediction of potential targets associated with microcephalia for the predicted ZIKV miRNAs

In order to comprehend the dynamics between viral miRNAs and their targets is extremely important to understand the complexity of biological regulation and virus-host interaction. *In silico* prediction of miRNA targets provides a suitable approach for identifying potential target sites based on their complete or partial complementarity with miRNAs. For these reasons, pairwise comparison of human gene transcripts of genes associated with microcephaly and miRNA-MD1, -MD13, -MD17 and -MD21 were performed by means of the use of the miRTar database [35]. The results of these studies are shown in Table 2. As it can be seen in this table, miRNA-MD1, and miRNA-MD17 hybridize with human gene transcripts of genes associated with micocephaly.

**Table 2.** Predicted ZIKV miRNA targets associated with microcephaly, identified by *in silico* analysis.

| Hairpin | Predicted miRNA | Targeted gene | Accession number | Protein description |
| --- | --- | --- | --- | --- |
| **MD1** | ZIKV-miRNA-MD1 | MCPH1 | AK022909 | Microcephalin 1 |
|  |  | WDR62 | BX647726 | WD repeat domain 62 |
|  |  | CDK5RAP2 | BK005504 | CDK5 regulatory subunit associated protein 2 |
|  |  | ASPM | AY367065 | Abnormal spindle microtubule assembly |
|  |  | CEP152 | AB020719 | Centrosomal protein 152kDa |
|  |  | ZNF335 | AK026157 | Zinc finger protein 335 |
|  |  | PHC1 | U89277 | Early development regulator 1 |
|  |  | CENPF | U30872 | Centromere protein F, 350/400ka (mitosin) |
| **MD17** | ZIKV-miRNA-MD17 | CENPF | U30872 | Centromere protein F, 350/400ka (mitosin) |

### Effective hybridization of predicted miRNAs and human gene transcript targets

In order to reconfirm effective hybridizations among the identified gene transcript targets and miRNA-MD1 and -MD17, we observed their hybridization patterns and calculated the minimum free energy for each hybridization. We used very stringent conditions, with a MFE of -30 kcal/mol as a cut-off value for miRNA-target pairing. The results of these studies are shown in Table 3.

**Table 3.**
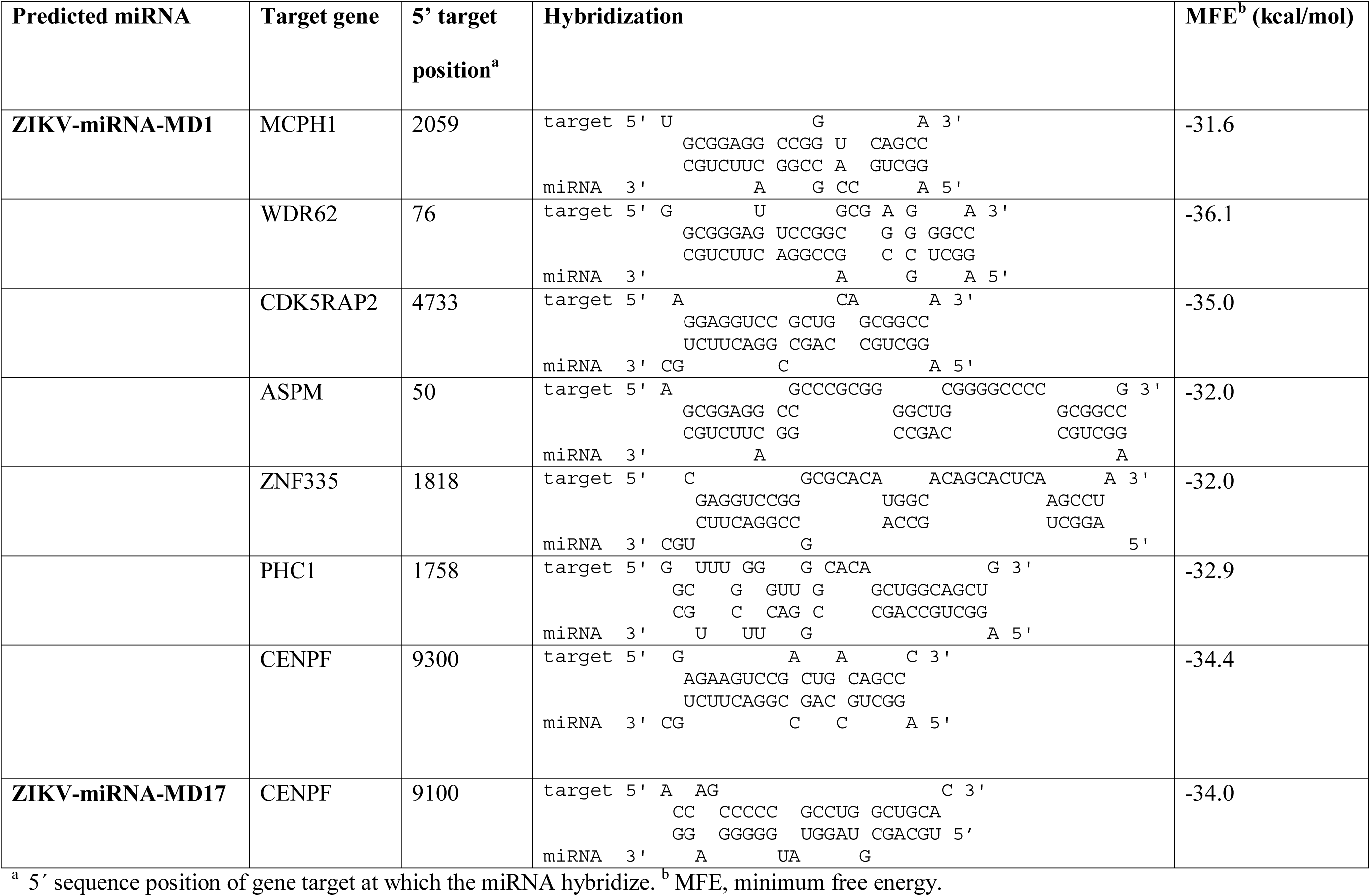
Effective hybridization of predicted ZIKV miRNA and transcript gene targets.

## Discussion

miRNAs play critical roles in many biological processes, such as cell growth, tissue differentiation, cell proliferation, embryonic development, cell proliferation, and apoptosis. As a consequence, their deregulation perturbs gene expression and can have pathological consequences [42].

Microcephaly is characterized by a reduced occipitofrontal circumference (OFC) of the head that is at least 4 standard deviations (SD) below the age- and sex-matched means and is caused by insufficient fetal brain development, which mostly affects the cerebral cortex [42]. Several genes have been mapped to date from various populations around the world in relation with microcephaly, including Microcephalin, WDR62, CDK5RAP2, CASC5, ASPM, CENPJ, STIL, CEP135, CEP152, ZNF335, PHC1, CDK6 and CENPF [43] (for gene names, accession numbers, protein description and role on brain development, see Supplementary Material Table 2).

Natal-RGN ZIKV strain was isolated from the brain tissue of a fetus diagnosed with microcephaly [10] and mouse models have provided significant evidence that ZIKV infection may cause brain defects by influencing brain cell development [12, 14]. Unfortunately, the mechanism by which ZIKV alters neurophysiological development remains unknown. A recent study revealed the miRNA coding capacity of ZIKV strains [29], which is in agreement with the results found in this work (see Table 1). In this study, using highly stringent conditions, four new miRNAs were found in the Natal-RGN ZIKV strain (Table 1). These miRNAs are situated in different ZIKV genome regions and are highly conserved among ZIKV strains (see Table 1 and Fig. 1). Moreover, two of these miRNA (ZIKV-miRNA-MD1 and MD17) were found to hybridize with target transcripts from genes previously shown to be associated with microcephaly (see Table 2). Futhermore, ZIKV-miRNA-MD1 effectively hybridize with MCPH1, WDR62, CDK5RAP2, ASPM, ZNF335 and CENPF (Table 2) even in stringent conditions (MFE > 30.0 kcal/mol) (Table 3), while ZIKV-miRNA-MD17 effective hybridizes with CENPF in the same conditions (Table 3). All these microcephaly associated genes have been found to be downregulated in an *in vivo* mice model of ZIKV infection, leading to disruption of neural development and microcephaly [14]. In this study, one of the predicted ZIKV miRNA (MD-21) is situated in the 3’untranslated region (3'UTR) of the ZIKV genome (Table 1). Flaviviruses are characterized by a relatively long and highly structured 3’UTR [44]. This region contains a number of stem-loops (SLs) and tertiary structures conserved among members of the genus *Flavivirus* [45], which makes it resistant to RNase degradation [46]. One of these SLs, the 3’SL located at the very end of the 3’UTR has been shown to be crucial for viral replication and interacts with a variety of proteins [47]. In order to gain insight into the location and possible functions of MD-21, the secondary structure of the 3’UTR of the consensus sequences obtained for ZIKV genomes included in these studies was established (Fig. 2). Interestingly, we found that the pre-miRNA-MD21 is identical to flaviviruses 3’SL and contains the short 5’-ACAG-3’ sequence in the top loop of the 3’SL, which is conserved among the members of this genus (see Fig. 2). Recent studies revealed the production of a miRNA (KUN-miR-1) from the 3’SL of the 3’UTR of WNV [27, 48]. Production of this miRNA was shown to be required for efficient WNV replication in mosquito cells. These studies also demonstrated that KUN-miR-1 functions *via* up-regulation of the expression of transcription factor GATA4, which in turn is required to facilitate WNV replication in mosquito cells [27]. In order to observe if GATA4 may be a target for ZIKV-miR-MD21, we computationally calculated that hybridization (Table 4). The result of this analysis revealed that ZIKV-miR-MD21 has the capability to hybridize *Aedes aegypti* GATA4 (Table 4).

**Fig 2.**
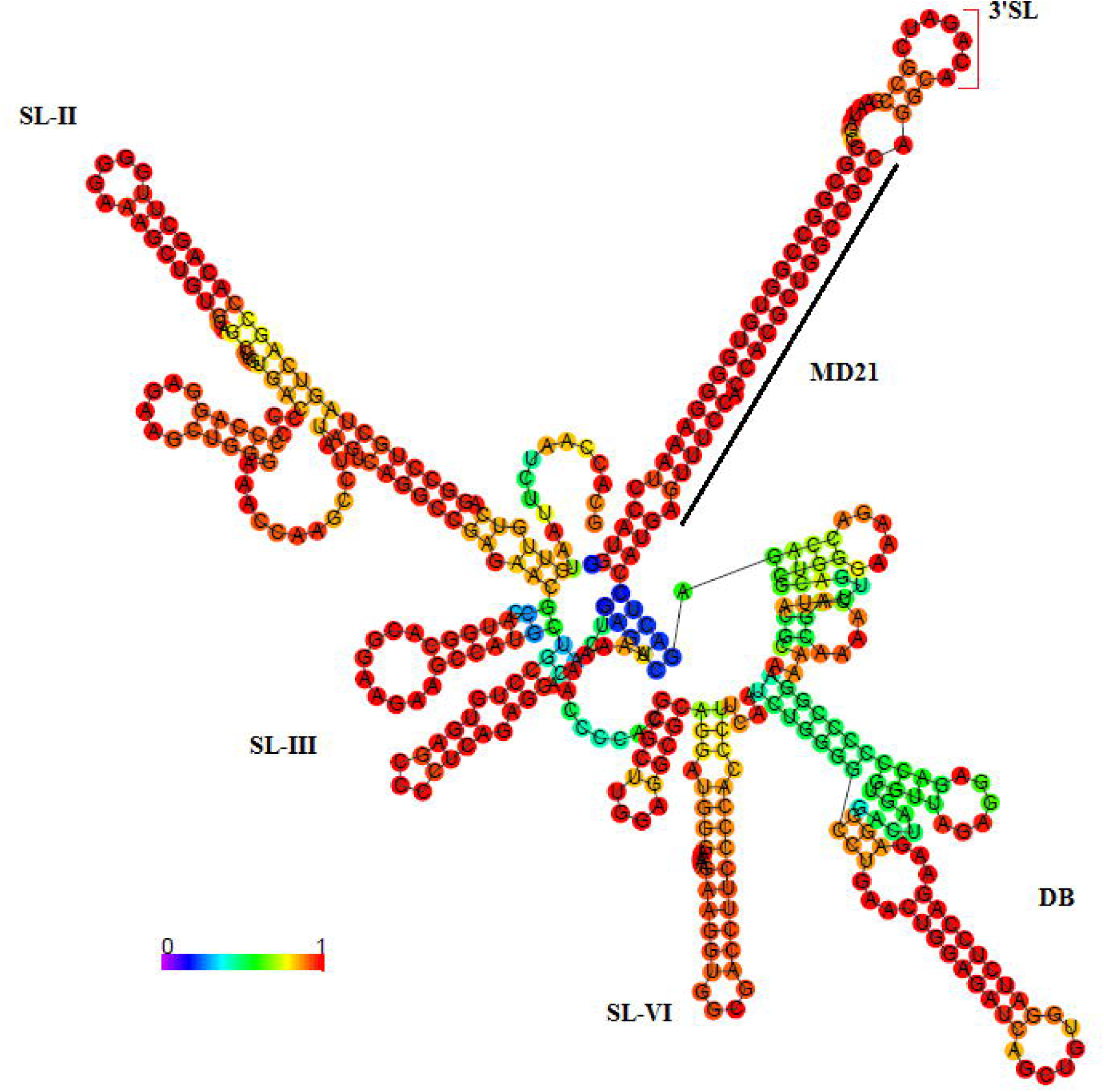
Predicted secondary structure of ZIKV 3’UTR. The secondary structure of 3’UTR consensus sequences obtained for the ZIKV strains included in these studies is shown. Stem-loops (SLs) names are indicated next to each SL. Flavivirus conserved 5’-ACAG-3’ sequence in the top loop of the 3’SL is indicated by a red bracket. Mature miRNA-MD21 sequences are indicated by a black line next to the sequences. Bar at the bottom of the structures denote base pair probabilities.

**Table 4.**
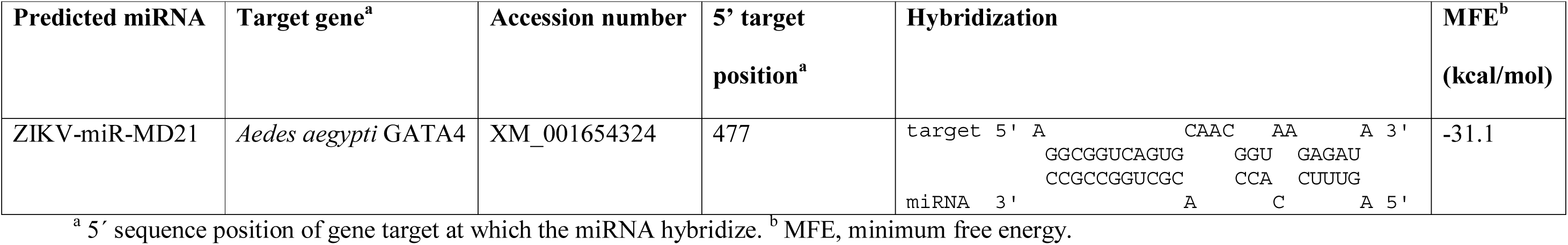
Effective hybridization of ZIKV-miR-MD21 and *Aedes aegypti* GATA4 gene.

Additionally, a recent study has reported another miRNA also produced from hairpin structures located in the 3’UTR of DENV-2 in infected mosquito cells [28].

Thus, there is increasing evidence suggesting the role of the 3’UTR of flaviviruses as a source of miRNAs. This is in agreement with the results of this work.

However, the details of the mechanisms for the generation of these miRNAs and their potential functions in virus replication in invertebrates hosts will require further investigation [44].

The results presented here suggest a possible role of miRNAs on the expression of human genes associated with microcephaly. Besides, they hint to candidate miRNAs that should be further confirmed by experimental analysis.

## Conclusions

The results of this work revealed the miRNA coding capacity of the ZIKV genome, in agreement with a recent report [29]. In this study, using highly stringent conditions, four new miRNAs were found in the Natal-RGN ZIKV genome, which was isolated from the brain tissue of a fetus diagnosed with microcephaly. These miRNAs are situated in different ZIKV genome regions and are highly conserved among ZIKV strains. Two of these miRNA (ZIKV-miRNA-MD1 and -MD17) were found to hybridize with target transcripts from genes previously shown to be associated with microcephaly. Moreover, effective hybridizations among ZIKV-miRNA-MD1 and -MD17 and these target genes can be observed even in highly stringent conditions (MFE > 30.0 kcal/mol). These results suggest a possible role of miRNAs on the expression of human genes associated with microcephaly and revealed candidate miRNAs that should be further confirmed by experimental analysis. Additionally, another new ZIKV miRNA, ZIKV-miRNA-MD21, was predicted in the 3’SL of the 3’UTR of the ZIKV genome, suggesting a potential role for the 3’UTR of flaviviruses as a source of miRNAs production.

## Acknowledgements

Authors acknowledge support by Agencia Nacional de Investigación e Innovación (ANII), PEDECIBA and Comisión Sectorial de Investigación Científica (CSIC), Grupos I+D, Universidad de la República, Uruguay.

## Supporting Information

**Supplementary Material Table 1. Origins of the ZIKV strains.**

**Supplementary Material Table 2. Human genes known to be associated with microcephaly.**

